# Decomposing the causes for niche differentiation between species using hypervolumes

**DOI:** 10.1101/485920

**Authors:** José Carlos Carvalho, Pedro Cardoso

## Abstract

1. Hutchinson’s n-dimensional hypervolume concept holds a central role across different fields of ecology and evolution. The question of the amount of hyper-volume overlap and differentiation between species is of great interest to understand the processes that drive niche dynamics, competitive interactions and, ultimately, community assembly.

2. A novel framework is proposed to decompose overall differentiation among hypervolumes into two distinct components: niche shifts and niche contraction / expansion processes. Niche shift corresponds to the replacement of space between the hypervolumes occupied by two species, whereas niche contraction / expansion processes correspond to net differences between the amount of space enclosed by each hypervolume.

3. Hypervolumes were constructed for two Darwin’ finches, *Geospiza conirostris* and *Geospiza magnirostris*, using intraspecific trait data from Genovesa Island, where they live in sympatry. Results showed that significant niche shifts, not niche contraction, occurred between these species. This means that *Geospiza conirostris* occupied a different niche space and not a reduced space on Genovesa.

4. The proposed framework allows disentangling different processes and understand the drivers of niche partitioning between coexisting species. Niche displacement due to competition can this way be separated from niche contraction due to specialization or expansion due to lower pressure on newly occupied ecological settings.

## 1 INTRODUCTION

The concept of species fundamental niche proposed by Hutchinson (1957) is at the roots of many ecological and evolutionary theories. Hutchinson formalized this concept as a multidimensional hypervolume, defined by a set of *n* independent variables that represent biologically relevant axes. According to his perspective, each point within this *n*-dimensional space corresponds to a possible state of the environment permitting the species to survive.

Hutchinsonian niches can be estimated by quantifying the functional trait hyper-volume occupied by the individuals of a given species. In essence, a set of functional traits, representing major axes of ecological strategies, are selected. Then, the hypervolume is constructed to estimate the morphospace occupied by the species. The hypervolume, representing the intraspecific trait variability, is commonly interpreted as the realized niche of the species (e.g. Pigot, Trisos, & Tobias, 2016).

The question of the amount of morphospace overlap and differentiation between species is of great interest to understand the processes that drive niche dynamics, competitive interactions and, ultimately, community assembly (e.g. Ricklefs & Cox, 1977; Stubs & Wilson, 2004; Kraft, Valencia, & Ackerly, 2008). The hypervolume overlap corresponds to the shared space between two species, whilst hypervolume differentiation corresponds to the sum of the unique fractions of space belonging to each species. Niche differentiation between species may be caused by two distinct processes: niche shifts and niche contraction (or expansion). Niche shifts are determined by the replacement of niche space between species. In this case, species change their niche position in order to occupy a different niche space. For example, morphological displacement of traits related to food acquisition have been interpreted as a strategy to reduce niche overlap and avoid competition (Huey, Pianka, Egan, & Coons, 1974). On the other hand, niche contraction (or expansion) processes are determined by differences in niche breadth (e.g. Pulliam, 1986). Niche contraction occurs when a species reduces its niche breadth. For example, when faced with more intense competition from another species, many organisms restrict their utilization of shared microhabitats and/or other resources (Pianka, 2000). This might give them advantage exploring these resources over other, generalist, species, as they adapt to explore them to the fullest. Niche expansion occurs when a species augments its niche breadth when presented with ecological opportunity (McCormack & Smith, 2008). Compelling examples of niche expansion are given by studies in islands where species were released from competition (e.g. Lister, 1976). Thus, to understand the factors that drive niche differentiation between species it is important to disentangle niche shifts from niche expansion (or contraction) processes.

In this paper, we propose a novel framework for partitioning hypervolumes via pairwise comparisons into niche shifts and niche contraction (or expansion). We present a case study to illustrate how these concepts can be applied to decompose the causes for niche differentiation between species.

## 2 MATERIAL AND METHODS

### 2.1 Hypervolume partitioning

To decompose hypervolume differentiation into different fractions, we use the set theory notation, in the manner of Hutchinson. Given a pairwise hypervolume system composed by hypervolumes *h*_*i*_ and *h_j_*, the total space occupied by them is given by their reunion (*h*_*i*_ ⋃ *h*_*j*_). Total pairwise hypervolume corresponds to the sum of hypervolume overlap given by the interception (*h*_*i*_ ⋂ *h*_*j*_) plus hypervolume differentiation, denoted by *h*_*i*_ Δ *h*_*j*_.

Thus, a pairwise hypervolume system can be partitioned according to the equation:

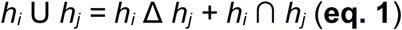

The differentiation between the hypervolumes *h*_*i*_ and *h*_*j*_ (*h*_*i*_ Δ *h*_*j*_) corresponds to the sum of their unique fractions:

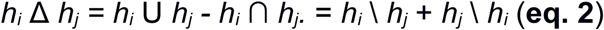

Total differentiation (*h*_*i*_ Δ *h*_*j*_) can occur as a consequence of two distinct processes: i) the differentiation that results from the replacement of space between hypervolumes; and, ii) differentiation that results from the net difference between the amounts of space enclosed by each hypervolume.

The net difference between both hypervolumes is given by |*h*_*i*_ \ *h*_*j*_ - *h*_*j*_ \ *h*_*i*_|. The replacement component can be obtained by subtracting the difference fraction from total differentiation, using equation 2:

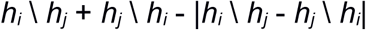

This expression can be written in the form:

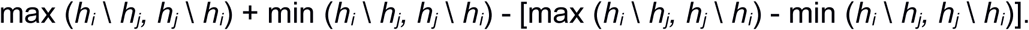

By simplifying the expression, we obtain the replacement component:

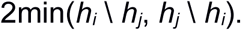

Thus, the total differentiation component can be additively partitioned in two components:

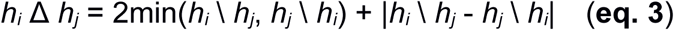

A total partitioning of *h*_*i*_ ⋃ *h*_*j*_ can be obtained by the equation (see Fig. 1):

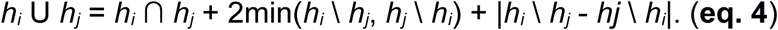

**Fig. 1.**
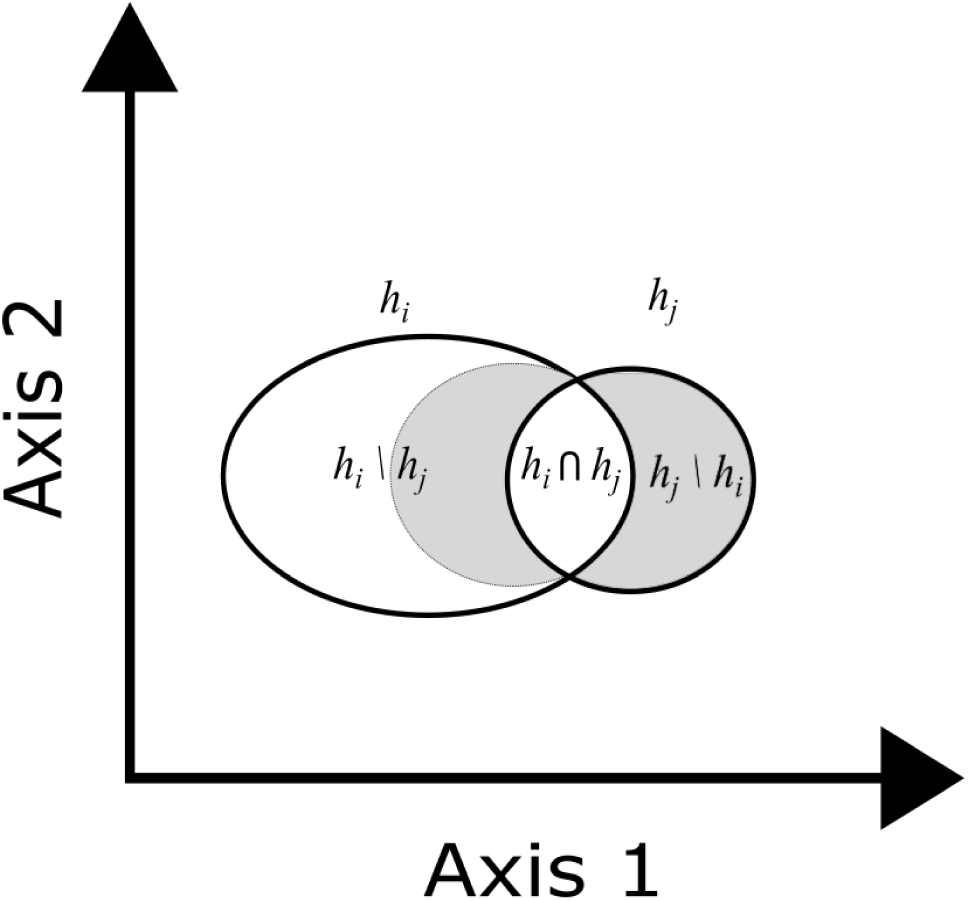
Diagram showing hypervolume partitioning in two dimensions. The overlap between hypervolumes *h*_*i*_ and *h*_*j*_ is given by the interception (*h*_*i*_ ⋂ *h_j_*) and the differentiation (*h*_*i*_ ∆ *h_j_*) is given by the sum of their unique fractions, *h*_*i*_ \ *h*_*j*_ and *h*_*j*_ \ *h_i_*. Note that: *h*_*i*_ ∆ *h*_*j*_ = 2min(*h*_*i*_ \ *h_j_*, *h*_*j*_ \ *h_i_*) + |*h*_*i*_ \ *h*_*j*_ - *h*_*j*_ \ *h_i_*|, where 2min(*h*_*i*_ \ *h_j_*, *h*_*j*_ \ *h_i_*) corresponds to the differentiation due to the replacement of space (grey shaded area) and |*h*_*i*_ \ *h*_*j*_ - *h*_*j*_ \ *h*_*i*_| refers to the net difference between both hypervolumes (white area in the *h*_*i*_ \ *h*_*j*_ space).

The terms in equations 3 and 4 can be scaled in relation to the total space occupied by the hypervolume pairwise system, thus obtaining the following equivalency:

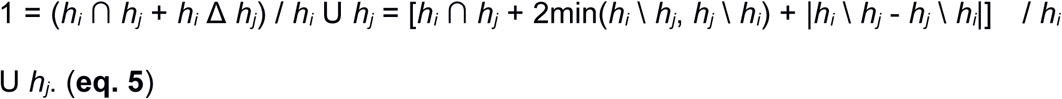

This equivalency can be summarized in the equation:

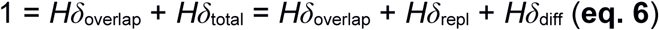

where, *Hδ*_overlap_ refers to the proportion of overlap between hypervolumes (similarity), *Hδ*_total_ refers to total differentiation between both hypervolumes, *Hδ*_repl_ corresponds to the proportion of differentiation that results from the replacement of space between hypervolumes and *Hδ*_diff_ is the proportion of differentiation that results from the net differences between the spaces occupied by both hypervolumes.

### 2.2 Assessing intraspecific trait variability and niche partitioning

Here, we advocate that intraspecific trait variability can be used to estimate niche parameters. The underlying assumption of a trait-based approach is that traits reflect species adaptations to the environment (Diaz & Cabido, 2001) and, hence, should be a useful tool to quantify species’ niches (Violle & Jiang, 2009). In other words, we are assuming that the morphospace occupied by each species is a surrogate for its realized niche.

Consider *N* individuals belonging to a given species *S* and measure a set of *T* traits for each individual. The matrix *N* × *T* can be used to construct the hyper-volume, which is assumed to describe the realized niche of the species. Therefore, the niche breadth of species *S* is given by the extent of the hypervolume.

Obviously, hypervolumes can be calculated for a set of species. The set operations can be applied to hypervolumes using eqs. 1-6 for each species pair. This allows obtaining a pairwise partitioning of the niche species pairs. Therefore, the terms in eq. 6 mean: *Hδ*_overlap_ = niche overlap between two species; *Hδ*_otal_ = total niche differentiation between two species; *Hδ*_repl_ = differentiation due to the replacement of niche space; *Hδ*_diff_ = differentiation due to differences of niche breadth between species. Thus, higher values of *Hδ*_repl_ are indicative of niche shifts between species, whereas higher values of *Hδ*_diff_ are indicative of niche contraction / expansion of one of the species in relation to the other (functional richness).

### 2.3 Incorporating different data types in hypervolume estimation

A limitation to the current hypervolume algorithms, is that they require datasets with continuous variables (Blonder, Lamanna, Violle, & Enquist, 2014; Blonder et al., 2018). We suggest that this limitation may be overcome by applying already available statistical methods to the trait matrix, from now on called **T** matrix, prior to hypervolume estimation.

Multivariate techniques, specially developed to deal with mixed data, may be used to ordinate the **T** matrix (e.g. Hill & Smith, 1976; Kiers, 1994). These ordination techniques are similar to a principal component analysis, allowing to extract several orthogonal axes (components) representing the variation of the **T** matrix (e.g. PCAMIX, Chavent, Kuentz-Simonet, Labenne, & Saracco, 2017). These axes can be interpreted in relation to the original variables (traits), by examining the matrix of eigenvectors in a similar way to a PCA. Then, the resulting axes are used as the new variables to compute hypervolumes (Fig. 2). If the aim is to include all the variation of the **T** matrix in the analyses, one should retain all the axes. However, most probably some of these components represent little variation and are difficult to interpret. Therefore, a more parsimonious strategy would be to select only those axes that provide a clearly interpretable result.

**Fig. 2.**
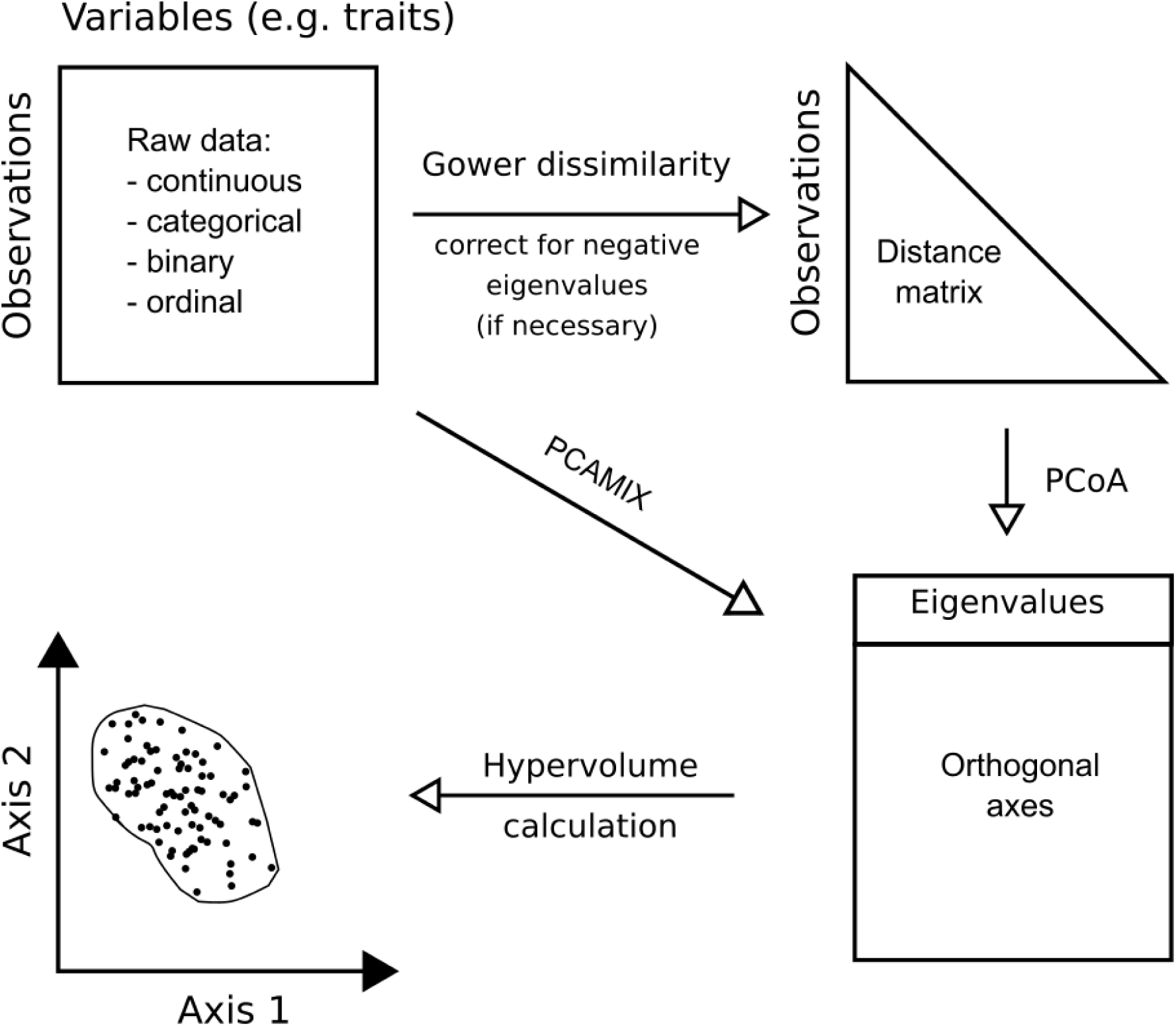
Diagram showing the construction of axes to be used in hypervolume estimation. The axes are obtained by ordination for mixed data types (PCAMIX; see Chavent et al. 2017) or a PCoA carried on a Gower dissimilarity matrix of individuals X traits (see Laliberté and Legendre, 2010). The eigenvalues correspond to the amount of variance explained by each axis.

The distance-based framework proposed by Laliberté & Legendre (2010) is an alternative to consider. Briefly, this approach is based on three steps: i) an appropriate distance measure (Gower dissimilarity measure; Gower,1971; Podani, 1999; Legendre & Legendre, 2012) is computed from the **T** matrix; ii) this distance matrix is analyzed through principal coordinate analysis (PCoA); and, (3) the resulting PCoA axes are used as the new variables to compute hypervolumes (Fig. 2). A comprehensive description of PCoA is given in Legendre & Legendre (2012).

Both PCAMIX and PCoA allow to extract orthogonal axes, which eliminates the potential correlation among trait variables, a pre-requisite of hypervolume methods. Independently of which technique is used to transform the **T** matrix, an additional point to consider is giving weights to trait variables, because a single categorical trait can be coded by several dummy variables. Also, prior weights can be given to trait variables, allowing to differentiate traits that contribute more to the biological performance of the species. Contrary to PCAMIX, the information of each trait is lost during the PCoA, because the **T** matrix is transformed into a dissimilarity matrix of individuals (or species), prior to the calculation of hypervolumes. Thus, the interpretation of these shapes is relative to the axes of the PCoA analysis and not in relation to the traits. Therefore, questions about which traits contribute more or less to a given hypervolume parameter are difficult to address.

### 2.4 Case study

We present a demonstration analysis of hypervolume partitioning for two species of Darwin’s finches: *Geospiza conirostris* and *Geospiza magnirostris*. Both species inhabit Island Genovesa, but only the former is present in Island Española. A well-known hypothesis for the two species is that competitive character displacement occurred on Genovesa and, therefore, they should have evolved to occupy dissimilar niches (Lack, 1945, 1947; Grant & Grant, 1982). We tested this hypothesis on Genovesa and Española Islands with morphometric data collected by Lack (1945, 1947): WingL, wing length; BeakH, Beak height; UbeakL, upper beak length and N-UBkL, nostril upper beak length. Data is available from Dryad Digital Repository: https://doi.org/10.5061/dryad.150. The original measurements (in mm) were log10-transformed prior to the construction of hypervolumes.

Hypervolumes for each species were calculated using a gaussian kernel density estimator with the hypervolume R package with default parameters (see Blonder et al., 2018 for details). Then, hypervolume decomposition (eqs. 1-6) was carried with the function *beta.volumes* of the BAT package (Cardoso, Rigal, & Carvalho, 2015). Hypervolumes are reported in units of SDs to the power of the number of trait dimensions used.

## 3 RESULTS

As hypothesized, hypervolumes calculated using morphometric data for *Geospiza conirostris* and *Geospiza magnirostris* were well differentiated on Genovesa Island (Fig. 3), the niche overlap between these species being null (*Hδ*_overlap_ = 0). By decomposing total differentiation (*Hδ*_total_ = 1) into replacement and differences between niche breadths, we found that such differentiation was mostly caused by niche replacement (*Hδ*_repl_ = 0.617) and not contraction/expansion, although the latter was also considerable (*Hδ*_diff_ = 0.383). By comparing the niche of *Geospiza conirostris* on Genovesa and Española, we found that they overlap slightly (*Hδ*_overlap_ = 0.242) and the differentiation between both islands was mainly due to the replacement of niche space (*Hδ*_repl_ = 0.711), not contraction/ expansion (*Hδ*_diff_ = 0.047). This result reinforces the interpretation that the presence of *Geospiza magnirostris* on Genovesa Island led to a significant shift (not contraction) of the niche of *Geospiza conirostris*.

**Figure 3.**
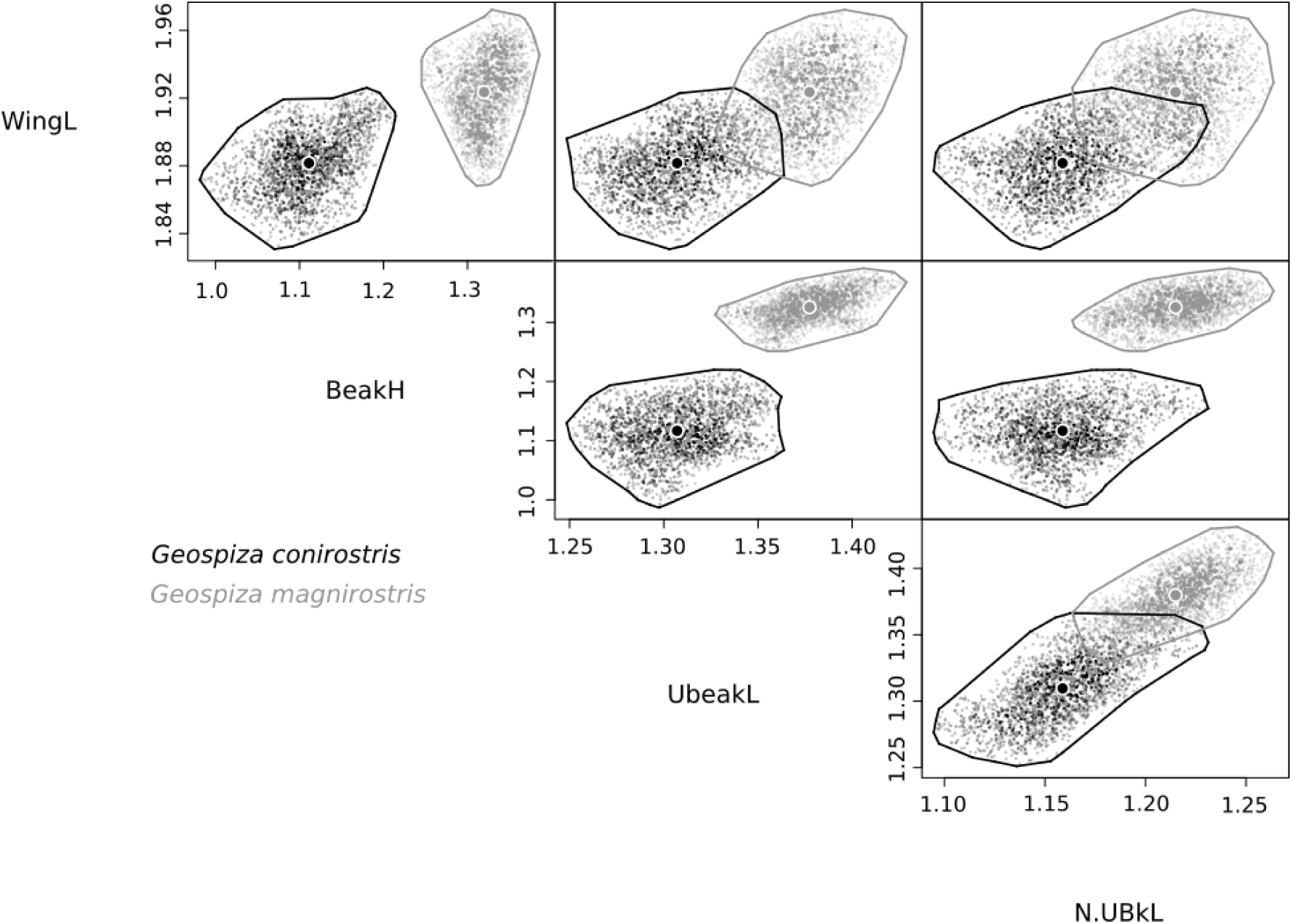
Niche partitioning for two species of Darwin’s finches *Geospiza conirostris* and *Geospiza magnirostris*. Hypervolumes shown as 2D projections for all combinations of trait axis (WingL - wing length; BeakH - Beak height; UbeakL - upper beak length and N-UBkL - nostril upper beak length) after log-transformation of the original measurements.

## 4 DISCUSSION

Hutchinson’s fundamental niche concept holds a central role across different fields of ecology and evolution. Conceptually, the species niche corresponds to an n-dimensional hypervolume enclosing the range of conditions under which the species can survive and reproduce. Niche overlap occurs when species use the same resources or strive under similar environmental conditions. In this regard, a crucial question is which processes determine the differentiation between species niches and, ultimately, determine species coexistence (MacArthur & Levins, 1967; Pianka, 1973). In this paper, we propose a novel framework to partitioning overall differentiation between hypervolumes into two distinct fractions: niche shifts corresponding to the differentiation that results from the replacement of space between hypervolumes and niche contraction / expansion, corresponding to the differentiation that results from the net differences between the space enclosed by hypervolumes.

We illustrate our framework with a classic dataset (Lack, 1945, 1947). Our results revealed that differentiation between *Geospiza conirostris* and *Geospiza magnirostris* occurred mostly by niche shifts processes and not by the niche contraction of *Geospiza conirostris* on Island Genovesa, where both species live in sympatry. This means that *Geospiza conirostris* occupied a different niche space and not a reduced space on Genovesa. These results are consistent with the hypothesis that the morphology of beaks was causally influenced by interspecific competition for food resources (Lack, 1947). This hypothesis received considerable support by analysing the food habits of these species (Grant & Grant 1982). These authors showed that *Geospiza magnirostris* feeds almost entirely on large-hard seeds, whilst *Geospiza conirostris* exploits *Opuntia* and arthropods on Genovesa. Moreover, on Española Island, where *Geospiza magnirostris* is absent, *Geospiza conirostris* consume large hard seeds. Therefore, it seems that the presence of *Geospiza magnirostris* induces a shift in the diet of *Geospiza conirostris* and, consequently, on its beak morphology.

The functional roles and the sensitivity to environmental changes of the vast majority of taxa are usually unknown (the so-called Hutchinsonian shortfall; Cardoso et al., 2011). Therefore, a trait-based quantification of species niche can provide a practical way to assess niche parameters for a potentially large number of species, making it possible to understand and predict species niche responses to environmental changes (Violla & Jiang, 2009; Blonder, 2018). Trait-based approaches have been used successfully to integrate functional ecology with community assembly and coexistence theories (e.g. McGill, Enquist, Weiher, & Westoby, 2006; Ackerly & Cornwell, 2007; Kraft, Valencia, & Ackerly, 2008). A major assumption of this approach is that trait dissimilarity is related to decreasing niche overlap among co-existent species (e.g. Stubs & Wilson, 2004), a rationale related to the limiting similarity principle (MacArthur & Levins, 1967). This premise assumes, implicitly, that the trait space is a good surrogate for the niche space. However, linking trait patterns to niche differentiation remains a challenge and, ultimately, depends on which traits were included in the analysis (D’Andrea & Ostling, 2016). Thus, an informed choice of traits should be made, based on their presumed adaptation to the species’ ecological niche. In conclusion, a novel framework was proposed to partition overall differentiation between hypervolumes into distinct fractions, replacement of space (niche shifts) and net differences between space amplitudes (niche contraction / expansion processes). The proposed framework allows disentangling different processes and understand the drivers of niche partitioning between coexisting species. Niche displacement due to competition can this way be separated from niche contraction due to specialization or expansion due to lower pressure on newly occupied ecological settings. We expect that the methods here presented will allow to address a wider range of niche- and trait-based questions than was possible to date.

## Acknowledgements

PC is supported by Kone Foundation with the project “Trait-based prediction of extinction risk”.

## Authors’ contributions

The authors contributed equally to the paper.

